# Mechanisms underlying melanoma invasion as a consequence of MLK3 loss

**DOI:** 10.1101/2021.12.10.472116

**Authors:** Henriette U. Balinda, Alanna Sedgwick, Crislyn D’Souza-Schorey

## Abstract

Invasive melanoma is an aggressive form of skin cancer with high incidence of mortality. The process of invasion is a crucial primary step in the metastatic cascade, yet the mechanisms involved are still under investigation. Here we document a critical role for MLK3 (MAP3K11) in the regulation of melanoma cell invasion. We report that cellular loss of MLK3 in melanoma cells promotes cell invasion. Knock down of MLK3 expression results in the hyperactivation of ERK, which is linked to the formation of a BRAF/Hsp90/Cdc37 protein complex. ERK hyperactivation leads to enhanced phosphorylation and inactivation of GSK3β and the stabilization of c-Jun and JNK activity. Blocking of ERK and JNK signaling as well as Hsp90 activity downstream of MLK3-silencing significantly reduces melanoma invasion. Furthermore, our studies show that ERK activation in the aforementioned context is coupled to MT1-MMP transcription as well as the TOM1L1-dependent localization of the membrane protease to invadopodia at the invasive front. These studies provide critical insight into the mechanisms that couple MLK3 loss with BRAF hyperactivation and its consequence on melanoma invasion.

## Introduction

Malignant melanoma accounts for about 10% of all skin cancers and is the deadliest type of skin cancer due to its highly metastatic nature and resistance to conventional chemotherapy and radiotherapy [1–3]. Patients with metastatic melanoma have traditionally had poor prognosis. Tumor metastasis is a multistep process wherein cancer cells invade through the extracellular matrix (ECM) and the surrounding tissues to disseminate to distant organs [4]. As such, the process of cell invasion is a prerequisite for tumor cells to metastasize. The majority of melanomas are associated with a deregulated oncogenic mitogen-activated protein kinase (MAPK) cascade involving Ras-Raf-MEK-ERK signaling [5], which is therapeutically most targeted in melanoma. The most prevalent and critical oncogenic mutation in cutaneous melanoma is in BRAF, a key serine-threonine kinase in the RAS-RAF-MEK-ERK (MAPK pathway) signaling pathway [5]. Approximately 90% of BRAF mutations are point mutations resulting in substitution of a valine with glutamic acid at amino acid 600 (V600E) [6]. Vemurafenib, a competitive kinase inhibitor with activity against BRAF(V600E), increases survival in patients although most patients become resistant to targeted therapy and relapse [7]. This underscores the need to better understand the molecular and cellular mechanisms underlying BRAF signaling in melanomas.

MAPK signaling pathways involve a three-step phospho-relay cascade, in which MAPKs are phosphorylated by MAPK kinases (MAP2Ks), that in turn are activated by MAPK kinase kinases (MAP3Ks). Mixed lineage kinase 3 (MLK3), also known as MAP3K11, is a member of the mixed lineage kinase (MLK) subfamily that belongs to a larger family of MAP3Ks. MLK3 is a serine/threonine kinase that is capable of activating several downstream MAPKs, including extracellular-signaling regulated kinase (ERK), c-Jun N-terminal kinase (JNK) and p38 in response to extracellular signals [8, 9]. Activation of MLK3 is associated with increased tumor cell invasion in several cancers, including ovarian, non-small cell lung and breast [10–12]. MLK3 activity has been implicated in vemurafenib resistance by directly activating MEK kinase leading to ERK activation and cell proliferation [13]. Other studies have shown that MLK3 functions as a scaffold to activate the RAF-MEK-ERK pathway [14, 15]. Here, using cell biological and biochemical approaches we have investigated the effects of MLK3 loss in melanoma invasion, wherein the function of the kinase remains to be fully understood. We show that cellular depletion of MLK3 increases melanoma invasion. This unexpected finding can be explained by the hyperactivation of ERK upon loss of MLK3 through the formation of a hyperactive BRAF/Hsp90/Cdc37 protein complex. Further, we demonstrate that MT1-MMP transcription and its localization mediated by TOM1L1 downstream of ERK activation is important for enhanced invasion resulting from MLK3 loss.

## Materials and Methods

### Antibodies

The following primary antibodies were used in this study: MLK3 antibody (Abcam), phospho-ERK, ERK, phopsho-JNK, JNK, beta-actin (Cell signaling Technologies), Raf-B and Cdc37 antibodies (Santa Cruz Technologies), phosphoserine (Millipore), alpha-tubulin (Sigma Aldrich) and TOM1L1 (Proteintech). Fluorescent secondary antibodies and rhodamine-phalloidin were purchased from Molecular Probes. HRP-conjugated secondary antibodies were purchased from Cell Signaling Technologies.

### Cell lines and cell culture

LOX and MDA-MB-231 cells were maintained in RPMI (GIBCO), supplemented with 2mM L-glutamine (Invitrogen), 10% FBS (HyClone), penicillin and streptomycin (Invitrogen). A375ma2 (ATCC) melanoma cells were maintained in high glucose DMEM (Sigma), supplemented with 2mM L-glutamine (Invitrogen), 10% FBS (HyClone), 1mM sodium pyruvate (VWR), penicillin and streptomycin (Invitrogen). Cells were grown in humidified incubator at 37°C with 5% CO_2_. For inhibitor experiments, cells were grown in the presence of CEP-1347 (1 μM) purchased from Tocris Bioscience, or PD98059 (25 μg/mL) purchased from Cayman Chemical, or JNK inhibitor II (also known as SP600125) (20 μM) purchased from Calbiochem, for 24 hours. 1μM of 17-AAG inhibitor was used for inhibition of Hsp90 activity.

### Lentiviral production and infections

MLK3-shRNAs in lentiviral vector pLKO.1-Puro plasmids were obtained from Sigma. Other plasmids used for lentivirus production were purchased from Addgene; pLKO.1 (Addgene plasmid 8453, originally obtained from Bob Weinberg), pMD2.G (Addgene plasmid 12259, originally obtained from Didier Trono), and psPAX2 (Addgene plasmid 12260, originally obtained from Didier Trono). To create the MLK3 shRNA-plasmid co-expressing GFP, EGFP was engineered into the pLKO.1 backbone for independent expression for independent expression of both proteins. Virus particles carrying MLK3 shRNA were generated and stored according to the manufacturer’s protocol. To carry out infection, cells were seeded at sub-confluence one day preceding infection. Virus was used at half strength, diluted with the normal culture media supplemented with 8 μg/mL polybrene. Culture media was changed 24 hours post infection and cells were incubated for an additional 48 hours prior to use. Infection efficiency was >90%.

### In vitro invasion assays

Gelatin degradation assays used here have been described previously [17, 18]. Briefly, 18 mm round glass coverslips (Fisherbrand) were coated with 2% gelatin (Sigma), which was conjugated to FITC or TRITC (Invitrogen). The gelatin coating was then cross-linked using 0.5% glutaraldehyde and excess aldehyde quenched with sodium borohydride (Sigma) and washed with excess 1X PBS. Cells were seeded onto coverslips and allowed to degrade matrix for 8 to 24 hours before being fixed and processed for immunofluorescence as described below to allow for the visualization of invasion/matrix degradation as dark spots or trails on a fluorescent background. Matrix degradation was quantified as previously described [17].

### Immunofluorescence staining and microscopy

For immunofluorescence staining, cells on gelatin-coated coverslips were fixed and stained as previously described [44], and mounted using Prolong Gold Antifade mounting media (Invitrogen). Images were obtained using either a Nikon A1R confocal microscope or a BioRad 1024 MRC confocal microscope. Image processing was performed using Image J (NIH).

### Immunoprecipitation assays

Immunoprecipitation assays were performed as described [44]. Briefly, cells were lysed in co-immunoprecipitation lysis buffer (Cell Signaling Technologies), supplemented with protease inhibitor cocktail and phosphatase inhibitor (Sigma) for 1 hour at 4°C with gentle rocking. Lysates were collected and the cell debris cleared by centrifugation. Equal amounts of total protein, determined by BCA assay (Pierce), were then pre-cleared using Protein A/G Plus beads (Santa Cruz). Cleared lysates were then incubated with primary antibody for 24 hrs at 4°C with gentle inversion. Beads were then washed 3 times with co-IP Lysis buffer. Washed beads were then resuspended in 2X SDS loading buffer (62.5 mM Tris-HCl, pH 6.8, 25% glycerol, 2% SDS and 0.01% bromophenol blue), heated to 100°C for 5 minutes and separated by SDS-PAGE.

### Western blotting assays

Western blotting assays were performed as described [22]. Briefly, cells were lysed in RIPA buffer (150 mM NaCl, 1 % IGEPAL CA-630, 0.5 % deoxycholic acid, 0.1 % SDS, 50 mM Tris-HCl pH 7.5) containing mammalian protease inhibitor cocktail (Sigma) and phosphatase inhibitor cocktail (Sigma) on ice. Protein concentrations were quantified using BCA assay (Thermo Scientific) and equal amounts of protein resolved on SDS-PAGE. Protein was transferred to PVDF membrane, and then blocked in a solution of 5 % milk in TBS-Tween (40 mL 1M Tris-HCl pH 7.4, 120 mL 5M NaCl, 2 mL Tween-20) at room temperature for 1 hour. Membranes were then probed with the indicated antibodies as per manufacturer’s recommendations.

### Statistics and reproducibility

Statistical analyses were performed using GraphPad Prism (version 6). Student’s t-test was performed when comparing two groups with data that appeared to be normally distributed with similar variances. When multiple groups were being analyzed and each group was compared to the control group and the data were normally distributed, we performed a one-way ANOVA with Dunnett’s multiple comparison test. All experiments were performed on at least 3 biological replicates under similar conditions.

## Results

### MLK3 silencing increases invasiveness of melanoma cells, but not breast cancer cells, through activation of ERK and JNK MAPKs

We initiated investigation of MLK3 activity on melanoma cell invasion by treating the invasive human melanoma cell line, LOX, with the pharmacological reagent, CEP-1347, known to interfere with activation of MLKs and downstream targets [16]. To examine invadopodia-mediated invasion, LOX cells treated with CEP-1347 or the vehicle DMSO as a control, were plated onto glass coverslips coated with fluorescently labeled, denatured collagen (gelatin), to monitor invadopodia-mediated invasion as described [17, 18]. CEP-1347 treatment rendered LOX cells significantly more invasive (Figures a-c). CEP-1347 treated cells were more flattened and spread with increased invadopodia-mediated matrix degradation that was observed as dark spots on the green fluorescent matrix relative to untreated controls (Figure 1a, c). Images taken along the z-axis (Figure 1b), showed increased cortactin containing invadopodia in CEP 1347-treated cells when compared to the control cells. CEP-1347 inhibits signaling pathways downstream of several members of the MLK family, so to confirm that the effect of CEP-1347 was MLK3-dependent, we examined cell invasion upon cellular depletion of MLK3 using small hairpin RNA (shRNA). Two shRNA sequences were used in this study and both were effective at silencing MLK3 expression (Figure 1d) in invasive melanoma cell lines, LOX and A375-MA2. When cells were infected with lentiviral vectors carrying shRNA against MLK3 (shMLK3), the cell lines show increased invasion following MLK3 silencing when compared to mock infected cells (Figure 1e-h). Intriguingly, expression of wild type MLK3 had no significant effect on melanoma cell invasion while expression of the kinase-dead mutant (K144R) exhibited a modest increase in cell invasion (Figure 1j). The results obtained were unexpected because previously MLK3 silencing, or inhibition, was shown to abrogate invasion in other tumor cell lines[19]. Hence, we examined the effect of depleting MLK3 in invasive breast cancer cells where inhibition of the enzyme correlates with reduced tumor cell invasion. In corroboration with previous findings, MLK3 shRNAs used here efficiently depleted MLK3 levels in MDA-MB-231 cells (Figure 2a) and markedly attenuated cell invasion (Figure 2b and c). MLK3 silencing in MDA-MB-231 cells also reduced the activation of the ERK and JNK kinases as previously reported (Figure 2d-f).

**Figure 1:**
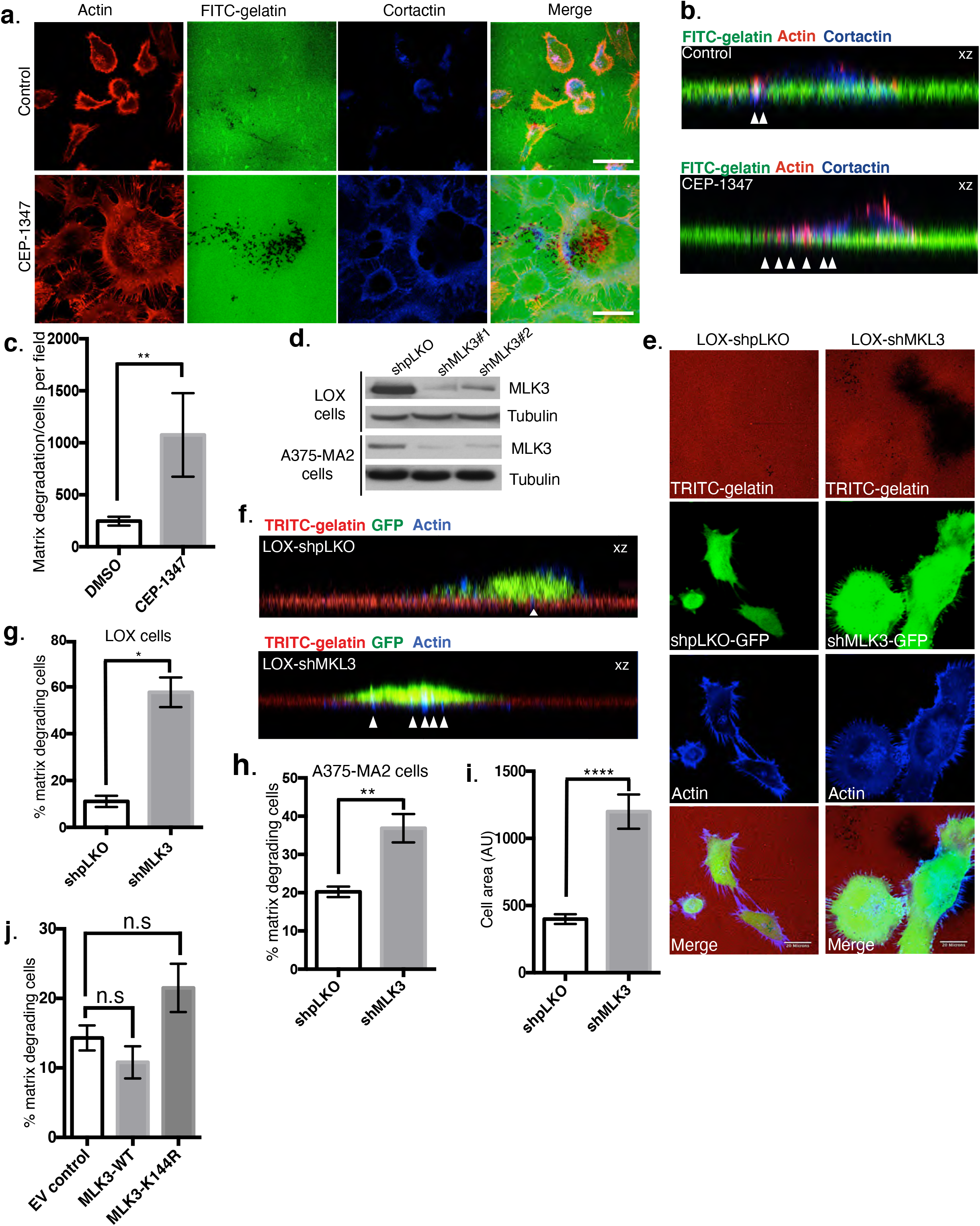
Cellular reduction of MLK3 increases matrix invasion of melanoma cells. **a-c)** LOX cells were grown on coverslips coated with FITC-gelatin matrix (green), treated with 2 μM CEP-1347, and incubated for 24 hours prior to fixing and staining for actin (red) and cortactin (blue). Cells were imaged along the x/y axis (a) or x/z axis (b). Quantification of matrix degradation is shown in (c). Error bars indicate standard error of the mean. **P-value = 0.0085. Matrix degradation was quantified by dividing pixels of dark spots by the number of cells in a field. **d)** LOX and A375-M2 cells were infected with lentivirus particles containing either the control shpLKO or shRNA against MLK3. Cell lysates were resolved by SDS-PAGE and probed using the antibodies indicated by Western blotting. **e-h)** LOX (e-g) or A375M2 (h) cells infected with lentiviral particles were plated on TRITC-labeled gelatin (red) and allowed to degrade matrix, fixed and labeled for actin (blue). Bar = 20 microns. Cells were imaged along the x/y axis (e) or x/z axis (f). g and h show percentage of cells degrading matrix. **i**) ImageJ was used to measure the cell surface area of 100 LOX cells with and without MLK3 depletion from 3 independent experiments. **j**) LOX cells were transfected with either the empty vector (EV) control, the MLK3-wild-type (WT) or the kinase dead mutant MLK3-K144R plasmids and then plated on FITC-labeled gelatin-coated coverslips and allowed to invade for 24 hours. Percentage of cells degrading matrix is shown. Error bars indicate standard error of the mean. *P-value <0.05; **P-value = 0.01; ****P-value < 0.0001.

**Figure 2:**
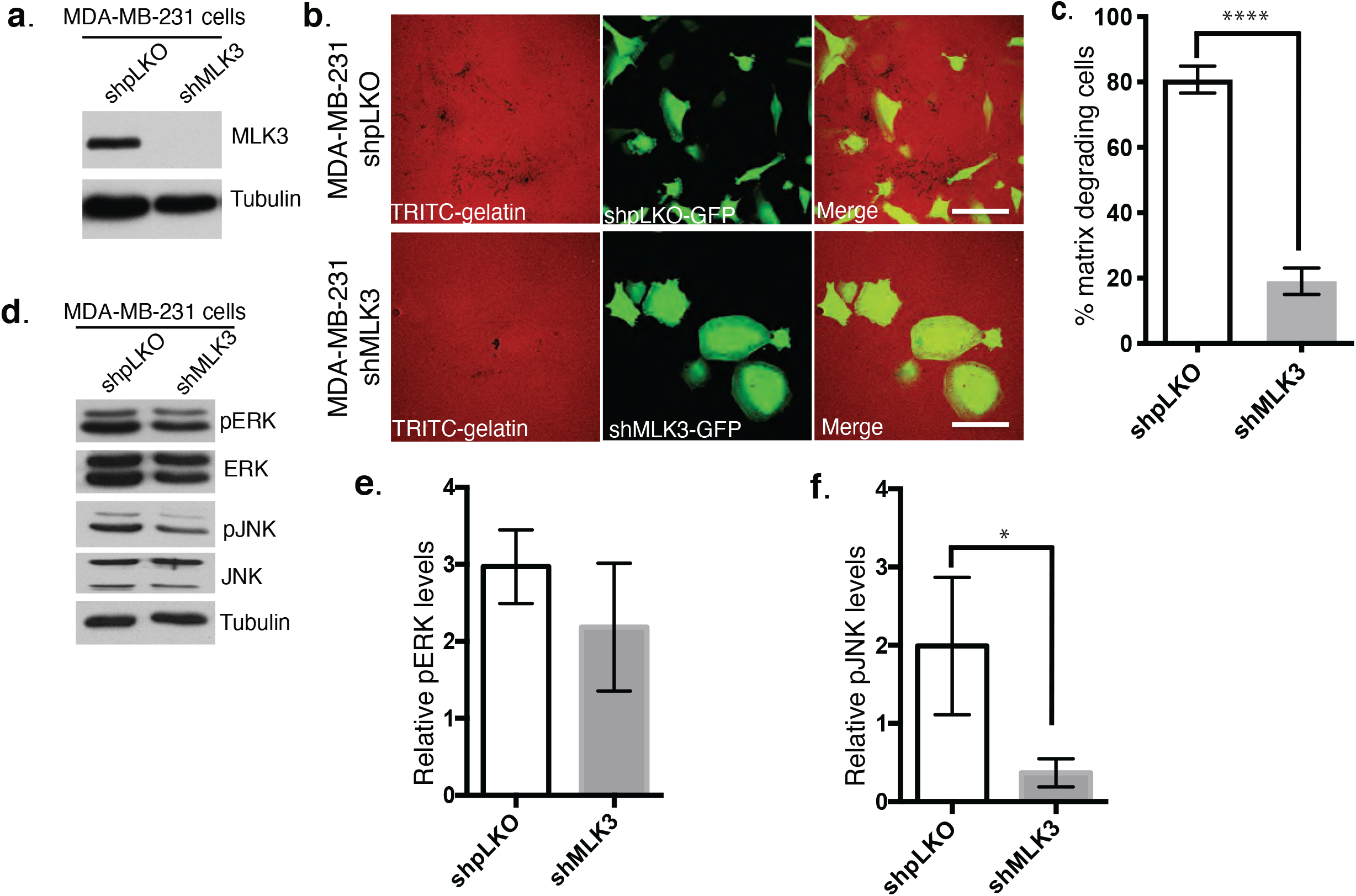
MLK3 silencing inhibits invasion in MDA-MB-231 breast tumor cells. **a)** Cells were infected with lentivirus particles encoding either the control vector or shRNA against MLK3. Cell lysates were resolved by SDS-PAGE and blotted using the antibodies indicated. **b)** MDA-MB-231 cells infected with lentiviral particles were plated on TRITC-labeled gelatin (red) and allowed to degrade matrix for 8 hours, fixed and viewed under confocal microscope. Bar = 20 microns. **c)** The percentage of cells degrading matrix is shown. **d)** Cells were infected with lentivirus encoding either shpLKO-GFP or shMLK3-GFP. Total protein lysates were then collected from the cells and analyzed by SDS-PAGE, and protein bands for pERK and pJNK (*p-value <0.05), were quantified using ImageJ.

Three major MAPKs, ERK, JNK and p38, are well characterized in the literature and MLK3 is known to modulate each of these MAPKs signaling cascades, [8, 20]. There is substantial data showing that the activation of ERK is essential to melanoma cell invasion [18, 21, 22]. To better understand the cellular mechanisms underlying the differences observed in MLK3 modulation of invasion in melanoma cells, relative to other invasive tumor cell lines such as MDA-MB-231, we investigated the activation status of JNK and ERK proteins by western blotting using the appropriate phospho-specific antibodies upon MLK3 silencing. We found that inhibition of MLK3 expression was accompanied by increased activation of ERK and JNK in LOX melanoma cells (Figure 3a-c).

**Figure 3:**
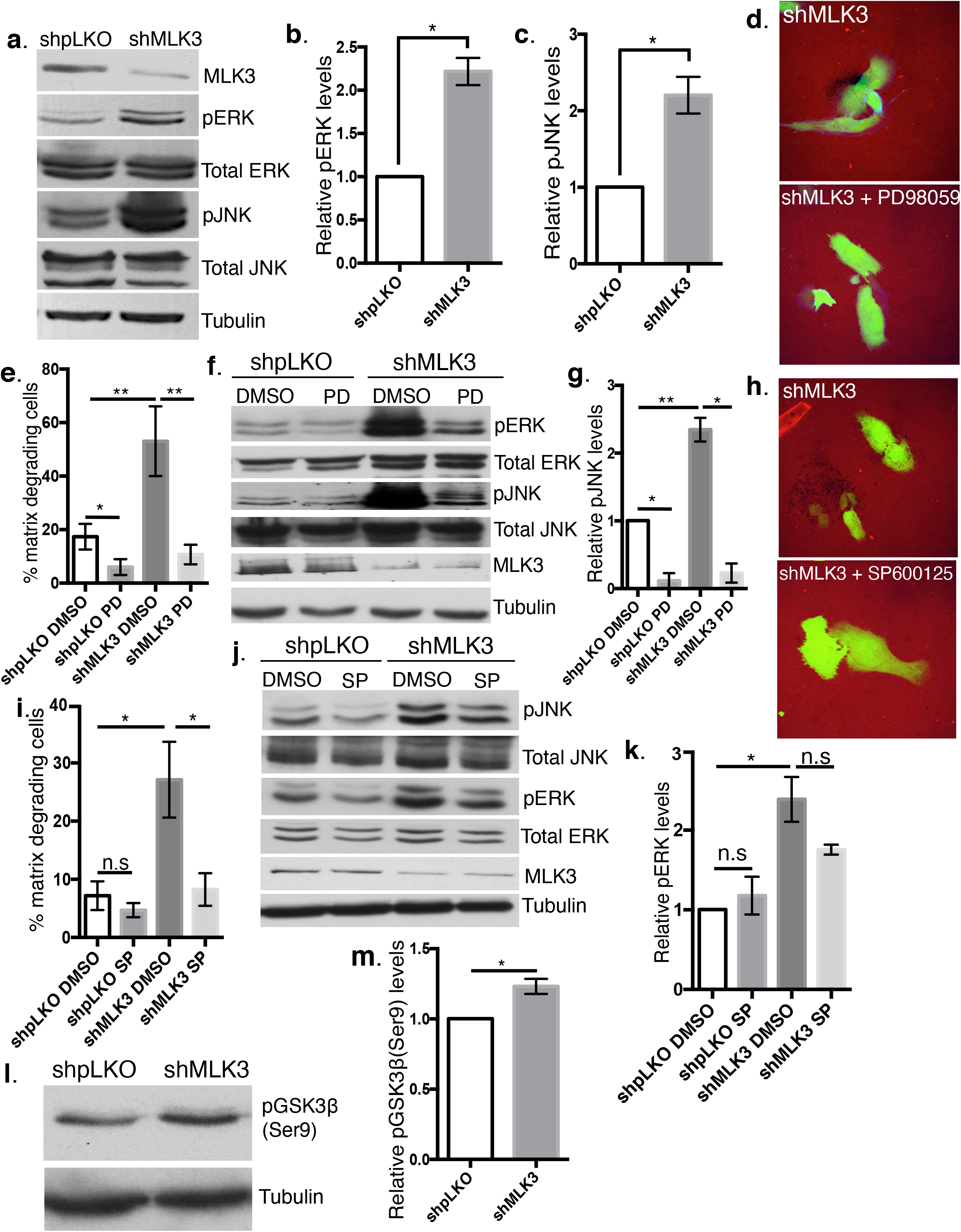
MLK3 silencing increases invasion through ERK and JNK activation. **a-c)** LOX cells were infected with lentivirus with either shpLKO-GFP or shMLK3-GFP plasmids for 72 hours. Protein lysates were then collected from the cells and analyzed by SDS-PAGE and probed using the antibodies indicated by Western blotting (a). (b) and (c) are bar graphs showing quantitation of proteins bands for pERK (p-value = 0.0163) and pJNK (p-value = 0.0152) using ImageJ. **d-e**) Infected LOX cells were plated on TRITC-labeled matrix (red) and treated with 50 μg/mL of PD98059, allowed to invade, fixed and labeled for actin (blue). Cells were counted and those with degradation spots underneath were noted and calculated as percentage of cells degrading matrix. Error bars indicate standard error of the mean. **f-g)** Protein lysates were collected from infected cells treated with or without 50 μg/mL PD98059 and analyzed by SDS-PAGE and probed using the antibodies indicated by Western blotting. Bands were quantitated using ImageJ. **h-i**) Infected cells with the appropriate shRNA were plated on TRITC-labeled matrix (red) and treated with 15 μM of SP600125 inhibitor and allowed to invade for 16 hours, fixed and labeled for actin (blue) as shown in (h). Matrix degrading cells were quantitated (i). **j-k)** Protein lysates were collected from infected cells treated with or without JNK inhibitor and analyzed by SDS-PAGE and probed using the antibodies indicated by Western blotting. Protein bands were quantitated using ImageJ. **l-m)** Total cell lysates were collected from cells infected with either shpLKO or shMLK3 and analyzed by western blotting for phosphorylated GSK3β on Serine 9. Error bars indicate standard error of the mean *p-value < 0.05. Bar = 20 microns.

### Inhibition of ERK and JNK hyperactivation downstream of MLK3 silencing reduces invasiveness in melanoma

To investigate the role of ERK activation in invasion downstream of MLK3 silencing, MLK3-depleted cells plated on fluorescently labeled matrix were treated with PD98059 the small molecule inhibitor of MEK, the kinase directly upstream of ERK, and allowed to invade the matrix. Inhibition of ERK downstream of MLK3 silencing in LOX melanoma cells leads to inhibition of invasion (Figure 3d and e). Moreover, ERK inhibition also decreased JNK activation (Figure 3f and g). This latter observation places ERK function upstream of JNK activation. The inhibition of JNK activity by treating cells with SP600125, a JNK inhibitor II, downstream of MLK3 silencing also resulted in the abrogation of increased invasion as a result of MLK3 silencing, but not in shpLKO (viral vector) control cells (Figure 3h and i). However, JNK inhibition resulted in an insignificant reversion of pERK levels, further supporting that the activation of ERK lies upstream of JNK in this context (Figure 3j and k).

Previous work has shown that constitutively active ERK can induce c-Jun stability via inhibitory phosphorylation of GSK3β at serine9, which in turn leads to enhanced JNK activity and a feed-forward mechanism of the JNK-Jun pathway [23]. Similarly, we observed that MLK3 silencing increased the levels of inhibitory phosphorylation of GSK3β-Serine 9, in line with the increase in JNK activation described above (Figure 3l and m). Taken together, these findings demonstrate that hyperactivation of ERK downstream of MLK3 silencing increases JNK activation through increased inactivation of GSK3β.

### MLK3 silencing leads to ERK hyperactivation through increased interaction between Hsp90/Cdc37 complex and BRAF

Two prior reports in the literature had direct bearing on these investigations. First, the Hsp90/Cdc37 complex protects MLK3 from binding to Hsp70 and its co-chaperone, an E3 ligase, and thus prevents the ubiquitination and degradation of MLK3 via the proteasome [24]. Second, a more recent study showed that BRAF^WT^ and BRAF^V600E^ localize in discrete protein complexes, and that BRAF^V600E^ isoform is found in a highly active protein complex with Hsp90/Cdc37 [25]. Given these reports, we hypothesized that silencing of MLK3 could promote the increased interaction between Hsp90/Cdc37 and BRAF leading to hyperactivation of ERK and increased invasion in melanoma. To test our hypothesis, we investigated the interaction between Hsp90/Cdc37 and BRAF upon MLK3 silencing using immunoprecipitation experiments. MLK3 silencing resulted in increased co-immunoprecipitation of endogenous Hsp90/Cdc37 and BRAF in both melanoma cell lines used in this study (Figure 4a-d). To investigate whether inhibition of Hsp90 downstream of MLK3 silencing would inhibit ERK hyperactivation, we utilized the Hsp90 inhibitor, 17-AAG, an analog of Geldanamycin. Inhibition of Hsp90 abrogated the increase in invasion downstream of MLK3 silencing (Figure 4e and f). Furthermore, 17-AAG treatment reduced ERK and JNK activity in MLK3-depleted LOX cells (Figure 4g-i). These findings provide a molecular basis for ERK hyperactivation (and increased cell invasion) downstream of MLK3 loss.

**Figure 4:**
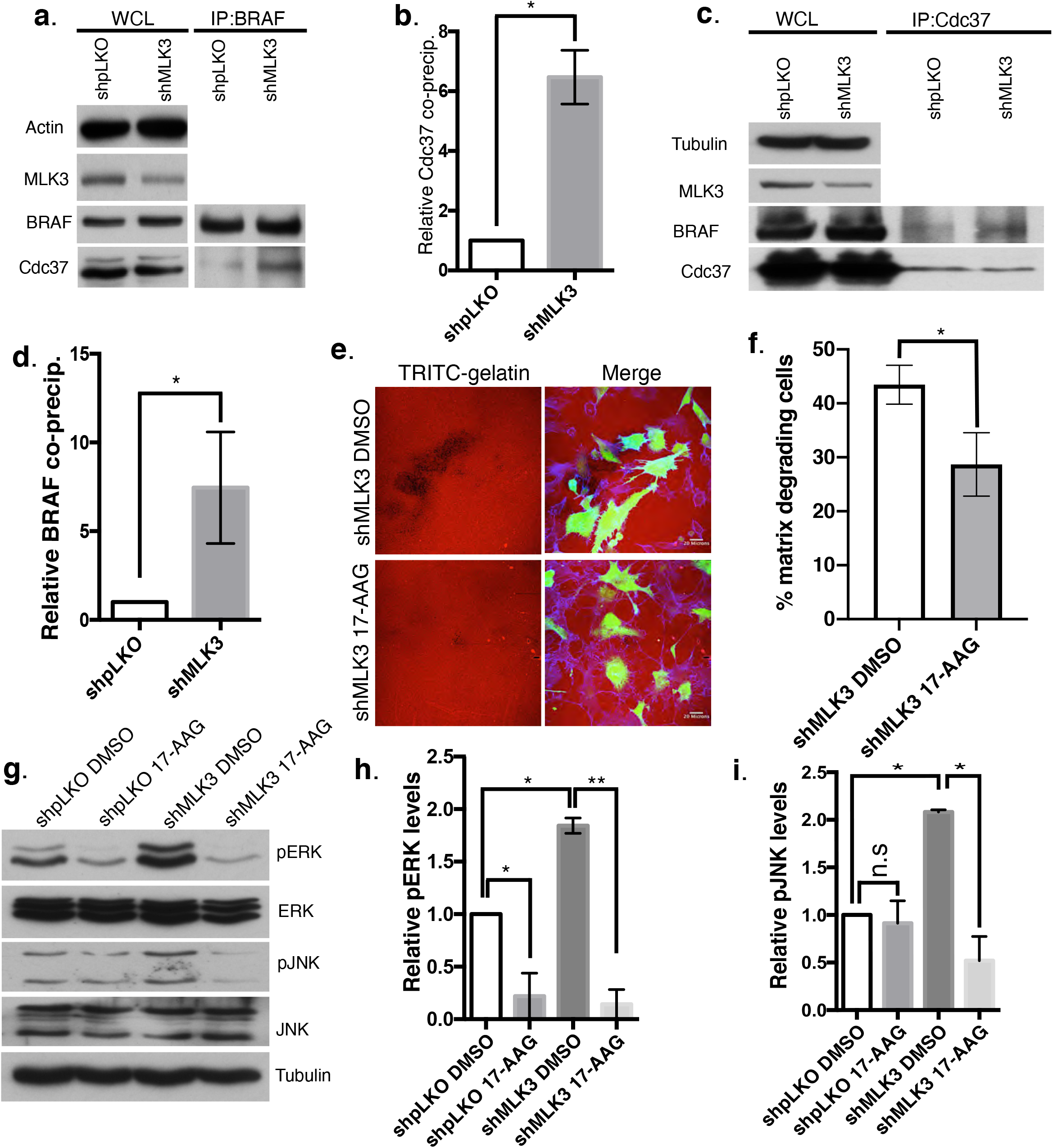
MLK3 silencing increases interaction between Hsp90/Cdc37 and BRAF. **a-d)** Lysates from control shpLKO- or shMLK3-infected cells were immunoprecipitated using monoclonal anti-Raf B (a) or anti-Cdc37 (c) antibodies. Whole cell lysates (WCL) and immunoprecipitates (IP) were resolved by SDS-PAGE, transferred to PVDF membrane and probed with the appropriate antibodies as indicated. (b) and (d) are graphs showing quantitation of the intensity of proteins bands in (a) and (c) respectively. **e-f)** LOX cells infected with either shpLKO control or shMLK3 were plated on TRITC-labeled (red) matrix, treated with DMSO as vehicle or 1 μM 17-AAG inhibitor, allowed to invade, fixed and stained for actin (blue) as shown in (e). Cells with degradation spots underneath were counted. The percentage of cells degrading matrix is shown (f). Error bars indicate standard error of the mean. Scale bar, 20 μM. **g-i)** Total cell lysates were resolved on SDS-PAGE, and probed with antibodies indicated using Western blotting. (h) and (i) are bar graphs showing quantitation of the blots in (g). *p-value<0.05, **p-value<0.01, ***p-value<0.005, ****p-value<0.001.

### MT1-MMP expression and localization govern cell invasiveness downstream of MLK3-silencing

Having observed that MLK3 silencing in invasive melanoma cells significantly increased ECM degradation as well as the hyperactivation of ERK and JNK, both of which are major activators of the AP-1 transcription factor [26], we hypothesized that the enhanced ECM degradation is likely mediated by increased levels of MT1-MMP, a master regulator of proteolytic degradation in melanoma. Consistent with this hypothesis, we observed that MLK3 silencing results in increased levels of both protein and mRNA levels of MT1-MMP (Figure 5a-c). We then investigated whether this increase in MT1-MMP was due to ERK /JNK activation. Treatment of cells with PD58059 or SP600125 inhibitors result in reduced MT1-MMP mRNA levels downstream of MLK3 silencing (Figure 5d). Notably however, as seen in Figure 5e and f, while we observed a consistent reduction in the pro-MT1-MMP protein in the presence of ERK and JNK inhibitors, the cellular levels of active MT1-MMP remain unchanged (discussed further below). Consistent with the above, pharmacological inhibition of MT1-MMP gelatinase activity using NSC405020, or of MMPs in general using the broad-spectrum inhibitor BB-94, inhibited the increased invasion induced by MLK3 silencing (Figure 5g and h).

**Figure 5:**
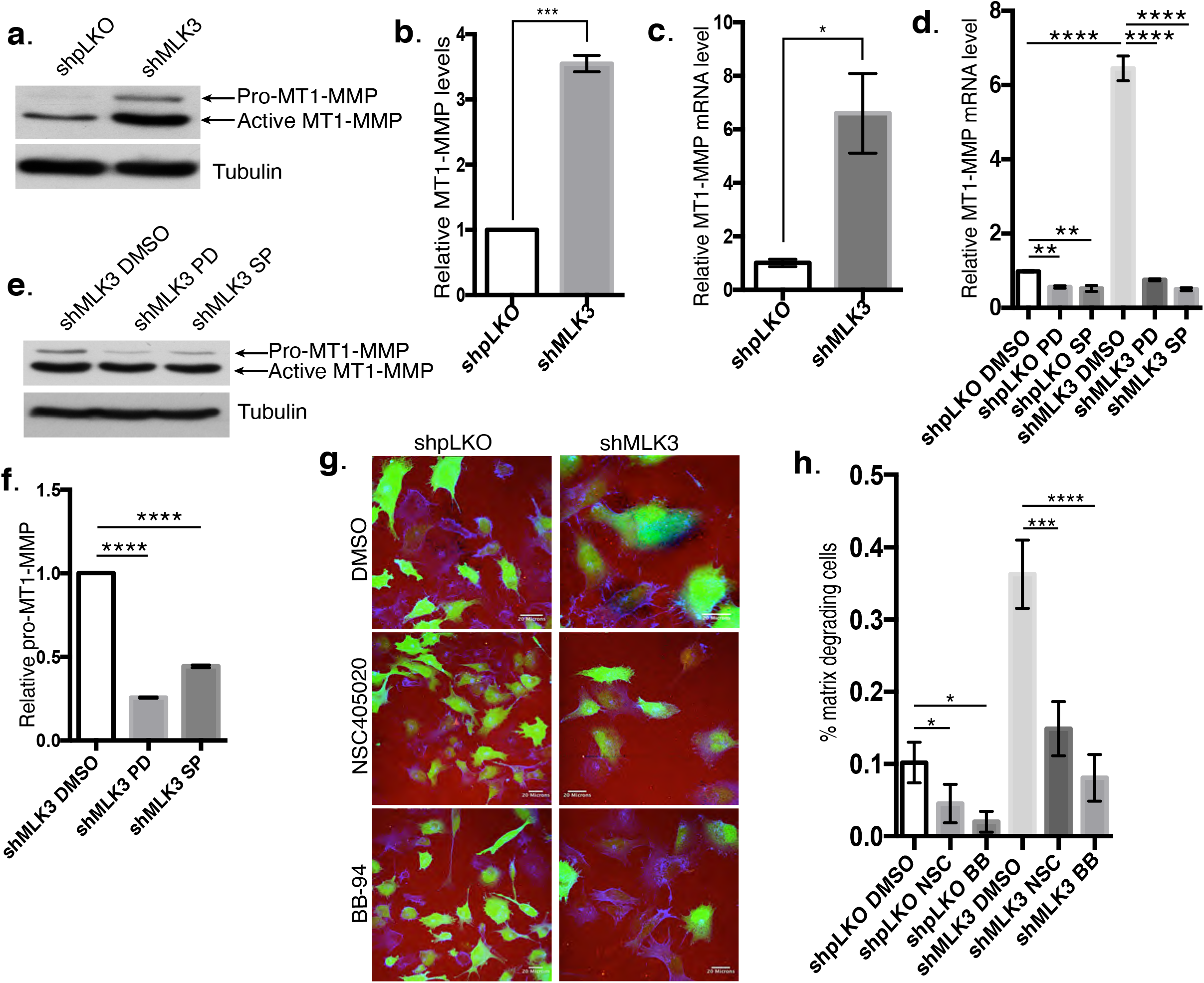
MLK3 silencing increases MT1-MMP levels. **a-b)** Total cell lysates were obtained from LOX cells expressing either shpLKO control or shMLK3, resolved on SDS-PAGE, and then probed by Western blotting with anti-MT1-MMP or anti-α-tubulin antibodies (a). (b) shows the quantitation of the blots in (a). **c-d)** mRNA was extracted from cells using the TRIZOL method, then reverse transcribed into cDNA and analyzed by RT-PCR. **e-f)** LOX cells expressing shMLK3 were treated with DMSO control or PD98059 or SP600125. Total cell lysates were obtained and resolved on SDS-PAGE, and probed using the antibodies indicated by Western blotting. Pro-MT1-MMP bands were quantitated using ImageJ (f). **g-h)** LOX cells expressing either shpLKO-GFP or shMLK3-GFP (green) were treated with DMSO or NSC405020 (100 μM) (MT1-MMP specific inhibitor) or BB-94 (10 nM) (pan MMPs inhibitor). The cells were then plated on TRITC-labeled gelatin (red), fixed and labeled for actin (blue) and images acquired using a confocal microscope. The percentage of degrading cells is shown in (h). The error bars indicate standard error. *p-value<0.05, ***p-value<0.005, ****p-value<0.001.

Given that ERK inhibition abrogates melanoma invasion, the high level of active MT1-MMP in LOX cells in the presence of the MEK inhibitor, PD98059, led us to question if localization of the active protease is perturbed in these cells. Consistent with previous findings, MT1-MMP localizes at sites of cell-ECM contact and especially at invadopodia (Figure 6a and b). However, this localization is lost upon treatment with PD98059 (Figure 6c). Instead, the majority of the MT1-MMP is redistributed to the dorsal surface not in contact with the matrix (Figure 6b, c). A recent study showed that the expression of TOM1L1 (Target of MYB1-like protein 1) regulates the localization of MT1-MMP at the cell-ECM contacts via a process that is dependent on the serine phosphorylation of TOM1L1 [27]. Thus, we examined TOM1LI phosphorylation by first immunoprecipitating cell proteins with anti-phosphoserine antibody with or without PD98059 inhibitor, and then probing for TOM1L1 levels by western blotting. Immunoprecipitation in reverse, with anti-TOM1L1 antibody and subsequent probing for phosphoserine, revealed the same result. Thus, ERK inhibition greatly reduces the levels of serine-phosphorylated TOM1L1 protein in shMLK3-depleted LOX cells (Figure 6e), resulting in altered localization of MT1-MMP.

**Figure 6:**
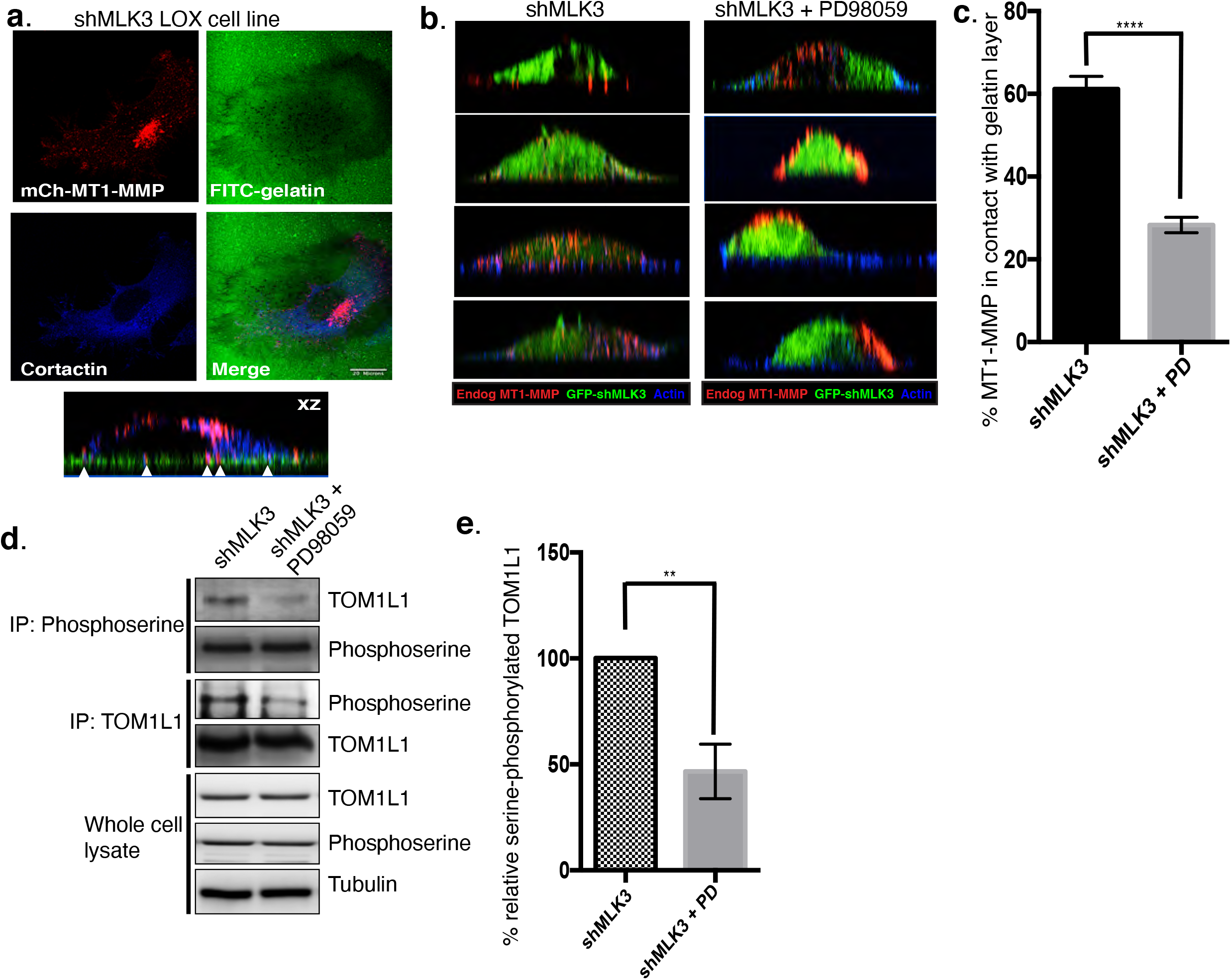
Inhibition of ERK activation downstream of MLK3 silencing inhibits localization MT1-MMP to invadopodia through reduced serine-phosphorylation of TOM1L1. **a)** LOX cells stably expressing shMLK3 were transfected with mCherry-MT1-MMP (red) plasmid assessed for invasion on FITC-labeled gelatin (green). Images along the x/y and x/z axes are shown. **b-c)** LOX cells were infected with lentivirus carrying GFP-shMLK3 plasmid, plated on unlabeled gelatin and either treated with PD98059 inhibitor or mock treated. Cells labeled for endogenous MT1-MMP and actin. MT1-MMP pixel intensity was measured using ImageJ. The amount of MT1-MMP in contact with gelatin was determined by the percentage of basal fluorescence of MT1-MMP divided by the total cell fluorescence. **d-e)** Lysates from shMLK3-infected cells or shMLK3-infected cells treated with PD98059 inhibitor were immunoprecipitated using anti-phosphoserine or TOM1L1 antibodies. Whole cell lysates and immunoprecipitates were resolved by SDS-PAGE and probed with the appropriate antibodies as indicated by Western blotting (d). Proteins bans were quantitated using ImageJ (e). The error bars indicate standard error. **p-value<0.01, ****p-value<0.001

## Discussion

This study provides new insights into MLK3 function in melanoma cell invasion. Our findings reveal a previously unappreciated role for MLK3 in the regulation of ERK hyperactivation in invasive melanoma via a pathway that involves the interaction of Hsp90/Cdc37 and oncogenic BRAF. Increased ERK activation phosphorylates GSK3β and enhances JNK activation and consequent expression of MT1-MMP. Further, ERK regulates localization of MT1-MMP to invadopodia via serine-phosphorylation of TOM1L1. Inhibition of ERK, JNK, Hsp90 or the membrane protease using small molecule inhibitors, abrogates increased melanoma invasiveness downstream of MLK3 silencing. Taken together, these findings have allowed us to propose a working model coupling MLK3 loss with BRAF hyperactivation and subsequent increase in melanoma invasion (Figure 7).

**Figure 7:**
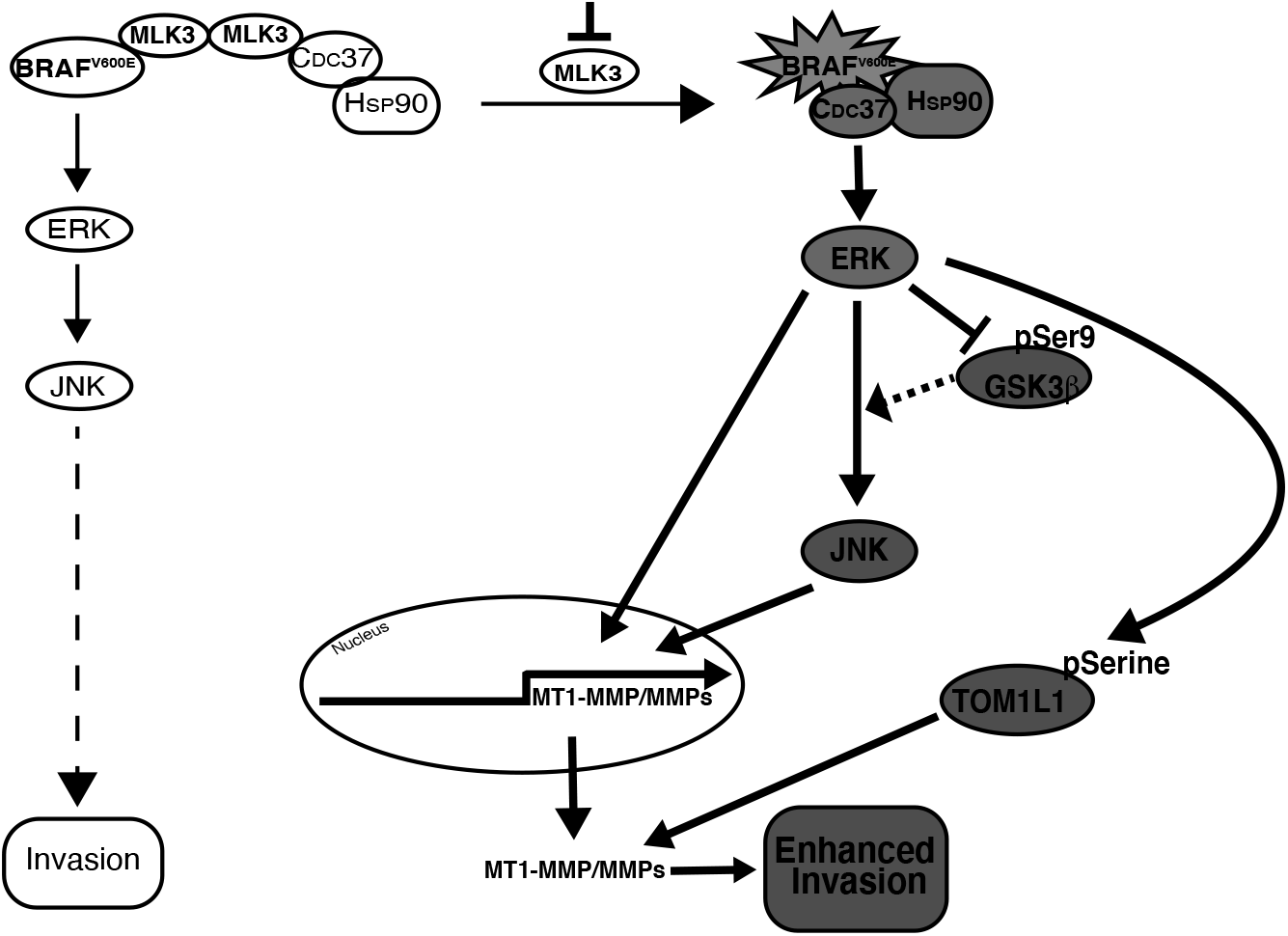
Working model for mechanisms involved in increased invasion downstream of MLK3. The data described in this study shows that inhibition of MLK3 results in increased melanoma invasion. Loss of MLK3 expression results in the hyperactivation of ERK, which is linked to the formation of a BRAF/Hsp90/Cdc37 protein complex. Enhanced ERK activation, leads to subsequent inactivation of GSK3β, and increased stability and activation of JNK. ERK activation also prompts transcription of MT1-MMP and the serine-phosphorylation of TOM1L1 leading to localization of MT1-MMP to invadopodia structures.

That MLK3 loss enhances melanoma cell invasion is in contrast to findings in breast, ovarian, small cell lung cancers and even in two melanoma cell lines, where loss of MLK3 function or expression, reduced tumor cell invasion [19, 28–31]. While our findings on the effect of MLK3 depletion in the invasive breast tumor cell line, MDA-MB-231 corroborate previous reports in the literature [19], we note that our findings in LOX and A375-MA2 melanoma cell lines are different from a previous study in Mel Ju and Mel Im melanoma cell lines which showed that depletion of MLK3 reduced invasion and migration capabilities [10]. The discrepancy of the latter study with the BRAF-associated mechanism proposed here is pertinent and will need further investigation. It is worth noting that utilized Boyden Chamber assays with Matrigel coated transwells to assess Mel Ju and Mel Im invasion. It is now appreciated that matrix proteolysis is an important process in regulating tumor cell invasion through crosslinked ECM barriers, and Matrigel, which is a reconstituted, non-crosslinked basement membrane, requires little if any proteolysis to breach [32, 33]. Further, melanoma-mediated invasion using the assays and cell lines described here requires MT1-MMP mediated proteolysis of the ECM.

The delivery of MT1-MMP to invadopodia is vital to ECM degradation and tumor invasion [34–37]. Here, MLK3 silencing is accompanied by an increase in MT1-MMP transcription and translation. Consistent with these observations, phospho-ERK and phospho-JNK are major regulators of transcription factors AP-1, which has been shown to regulate MT1-MMP expression [26]. Moreover, high levels of ERK activation, as observed here, has been shown to block the autodegradation of MT1-MMP and increase the stability of MT1-MMP through the formation of a complex with TIMP-2 and MMP-2[38, 39]. Blocking ERK or JNK activity downstream of MLK3 silencing decreased MT-MMP transcription but intriguingly, had no effect on the levels of active MT1-MMP. Rather, the inhibition of ERK downstream of MLK3-silencing resulted in a redistribution of MT1-MMP away from cell-ECM contact sites, a step dependent on the phosphorylation of TOM1L1. Also in line with our findings, available phospho-proteomic data (www.phosphosite.org) indicate that BRAF and MKK1/2 inhibitors modulate phosphorylation of TOM1L1 on serine 321 and 323 [40]. TOM1L1 was originally shown to aid in the translocation of MT1-MMP to invadopodia structures in ERBB2-driven invasion in breast tumor cells [27]. Thus, despite the differences in upstream signaling in different tumor cell types, there is remarkable similarity in the downstream mechanisms and target molecules involved.

Approximately 60% of melanoma tumors harbor activating BRAF mutations. The most frequently occurring BRAF mutation, BRAF(V600E)[6], is also found in the LOX and A375M2 cell lines used in this study [41]. Aberrant activation of the MEK/ERK pathway downstream of oncogenic BRAF in metastatic melanoma has been well established [42]. Although there is a growing body of evidence that ERK activation is important for the highly metastatic behavior of melanoma, much still remains to be investigated regarding ERK activation during melanoma progression. MLK3 has been reported to play a role in mitogen activation of BRAF and to function as a scaffold enabling ERK activation [8, 14]. As described here, we observe a significant increase in interaction between BRAF and Hsp90/Cdc37 upon MLK3 silencing leading to ERK hyperactivation. In addition, the data shows that inhibition of Hsp90 using 17-AAG, an analog of geldanamycin, blocks ERK activation and invasion downstream of MLK3 silencing. Thus, modulation of both MLK3 and Hsp90 provide prospects for therapeutic intervention in combination with current strategies of BRAF inhibition. Inhibition of BRAF^V600E^ by vemurafenib or dabrafenib is clinically important however, but almost invariably, resistance arises within a short time period [43]. All of the above underscore the need to better understand context-dependent regulation of mutant BRAF regulation and signaling at the molecular level.

